# Neural signatures of loss of consciousness and its recovery by thalamic stimulation

**DOI:** 10.1101/806687

**Authors:** Jacob A. Donoghue, André M. Bastos, Jorge Yanar, Simon Kornblith, Meredith Mahnke, Emery N. Brown, Earl K. Miller

## Abstract

We know that general anesthesia produces unconsciousness but not quite how. We recorded neural activity from the frontal, parietal, and temporal cortices and thalamus while maintaining unconsciousness in non-human primates (NHPs) with propofol. Unconsciousness was marked by slow frequency (∼1 Hz) oscillations in local field potentials, entraining local spiking to Up states alternating with Down states of little spiking, and decreased higher frequency (>4 Hz) coherence. The thalamus contributed to cortical rhythms. Its stimulation “awakened” anesthetized NHPs and reversed the electrophysiologic features of unconsciousness. Unconsciousness thus resulted from slow frequency hypersynchrony and loss of high-frequency dynamics, partly mediated by the thalamus, that disrupts cortical communication/integration.

---

Unconsciousness is general anesthesia’s most intriguing feature^1^. In humans, unconsciousness by propofol—a widely used anesthetic—is linked to structured neurophysiological dynamics: decreases in higher frequency (>15 Hz) but increases in frontal alpha (8 to 12 Hz)^2–5^ and slow frequency (0.1 to 1 Hz) oscillations that phase-lock neuronal spiking^6^ and loss of fronto-parietal connectivity.^7^ Nevertheless, the details remain unknown. These dynamics have been investigated using extracranial measures (EEG, fMRI) with limited spatial or temporal specificity or microelectrode recordings with limited spatial coverage in patients. They have yet to be studied at high spatial and temporal resolutions simultaneously. Moreover, previous studies have not included the thalamus, a critical nexus that regulates cortical activity^8,9^.

A vascular access port implanted subcutaneously in two non-human primates (NHPs) allowed us to infuse propofol to transition them from awake through loss of consciousness (LOC) and recovery of consciousness (ROC) without interruption of neurophysiological recording (and without introduction of other anesthetics or other confounding factors, see Methods, Figure 1A). We started with a high dose for 15 minutes (0.28-0.58 mg/kg/min, adjusted per individual animal). After LOC (indicated by cessation of airpuff-evoked eyeblinks, reduction in heart rate, pupil dilation, and decreased muscle tone), infusion was lowered to a holding dose (0.14-0.23 mg/kg/min). After infusion ceased, ROC followed after ∼60 sec. We recorded spikes and local field potentials (LFPs) from 64-channel “Utah” arrays in frontal cortex (8A, PFC), posterior parietal (PPC, 7A/B), and auditory/temporal (Superior Temporal Gyrus, STG) cortex and LFPs from multiple-contact stimulating electrodes (two in each hemisphere) in frontal thalamic nuclei (intralaminar nuclei, ILN, and mediodorsal, MD, Figure 1B).

**Figure 1.**
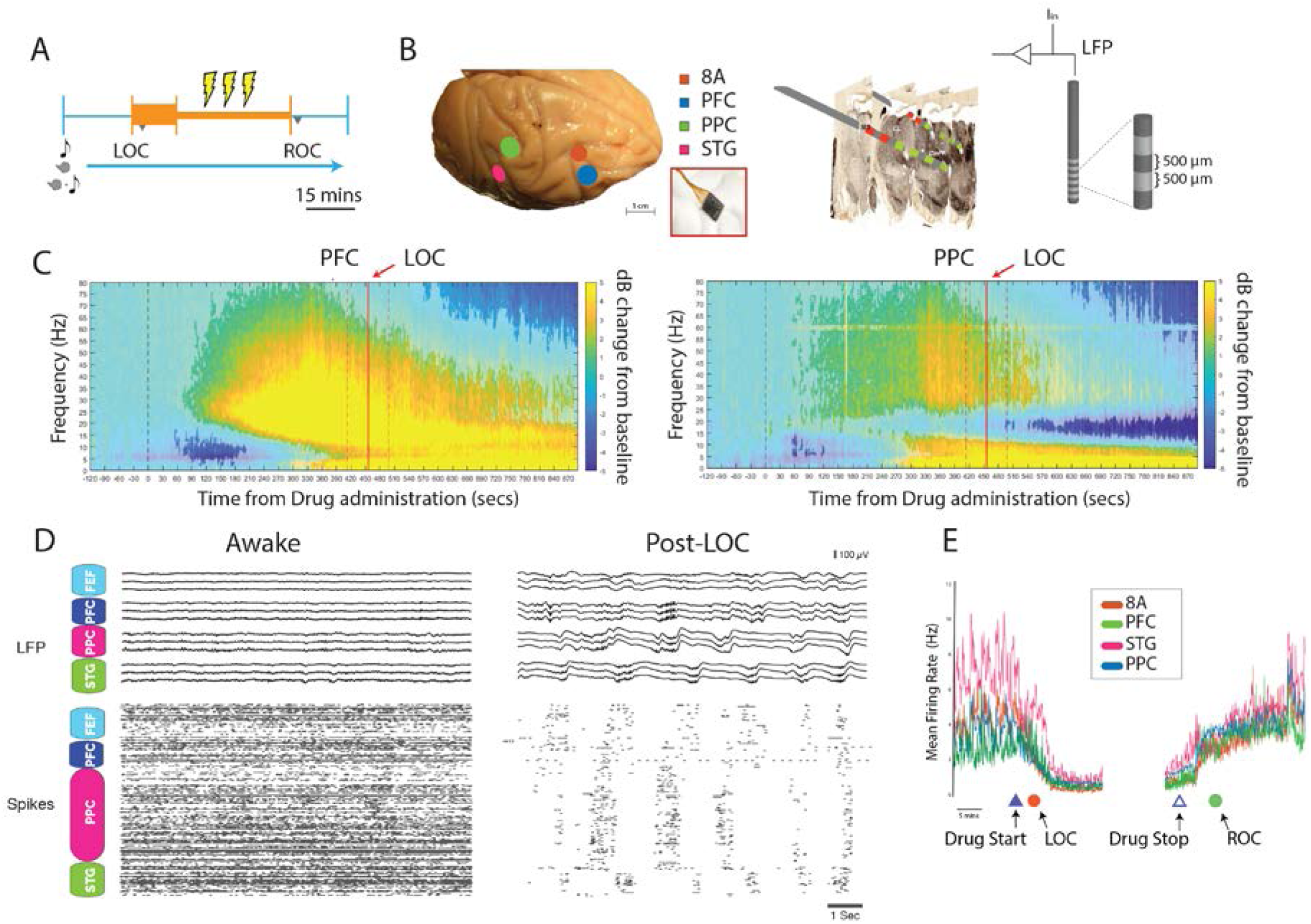
Paradigm, techniques and neurophysiological signatures of LOC. A. There was a 15-minute induction propofol infusion (fast rate, thick orange bar). After LOC, infusion switched to a halved rate of infusion for maintenance of LOC (narrow orange bar). 30-second thalamic stimulation (yellow bolts) occurred during the maintenance phase. LOC: Loss of consciousness, ROC: Recovery of consciousness. B. (left) Location of 64-channel chronic recording array. PFC: ventrolateral prefrontal cortex; 8A: caudal lateral PFC; PPC: posterior parietal cortex; STG: superior temporal gyrus (right) Central thalamic recording locations/stimulations with reconstruction of the histological sections. MD: mediodorsal nucleus of the thalamus, the CL: Central lateral nucleus and CmPf: centromedian/parafasicular complex (intralaminar nuclei). Green: example of sites of effective stimulation. Red: ineffective sites. Schematic of multi-contact thalamic electrodes. C. (left) Changes in power relative to pre-drug baseline in area 8A in decibels (positive numbers indicate enhanced, negative numbers decreased power). Significant changes are shown by more saturated colors. Solid vertical bar indicates average time of LOC, dashed lines the SEM. (right) Same as A but for posterior parietal cortex (PPC). D (left upper panel) Example LFP traces from the four cortical arrays during awake state. (left lower panel) Example spike rasters. (right panels) same as D, but for post-LOC. Note the synchronous LFP and spiking in the SF (∼0.5-1.5 Hz) band. E. Mean spike rate across all units in all recorded areas from an example session.

We compared LFP power during the pre-drug Awake state (−120 to 0 seconds pre drug-onset) to every timepoint after drug administration (non-parametric cluster-based randomizations, corrected for multiple comparisons; all effects p<0.01^10^). Around 60-75 seconds post-drug administration (pre-LOC), theta/alpha (4-15 Hz) power decreased in all areas (e.g., Fig 1C, Supp Figs 1, 2). Gamma (>35 Hz) and high-beta power (peak frequency ∼25-28 Hz) temporarily increased in frontal cortex (8A and PFC) and thalamus. High beta (not seen in PPC, STG; Supp Fig 1) then slowly ramped down to 15 Hz around LOC (∼10 minutes post drug-administration). Just before LOC (about 300 seconds post-drug administration) lower frequency (<4 Hz) power increased in all areas, peaking after LOC (at 455s +/-43s, two SEM across sessions, N=15; Supp Figs 1, 2). PPC alone then showed a drop in lower beta power (15-20 Hz) shortly after LOC. Similar dynamics are seen in human EEG^4,11^. Below, we compare Awake to post-LOC.

**Figure 2.**
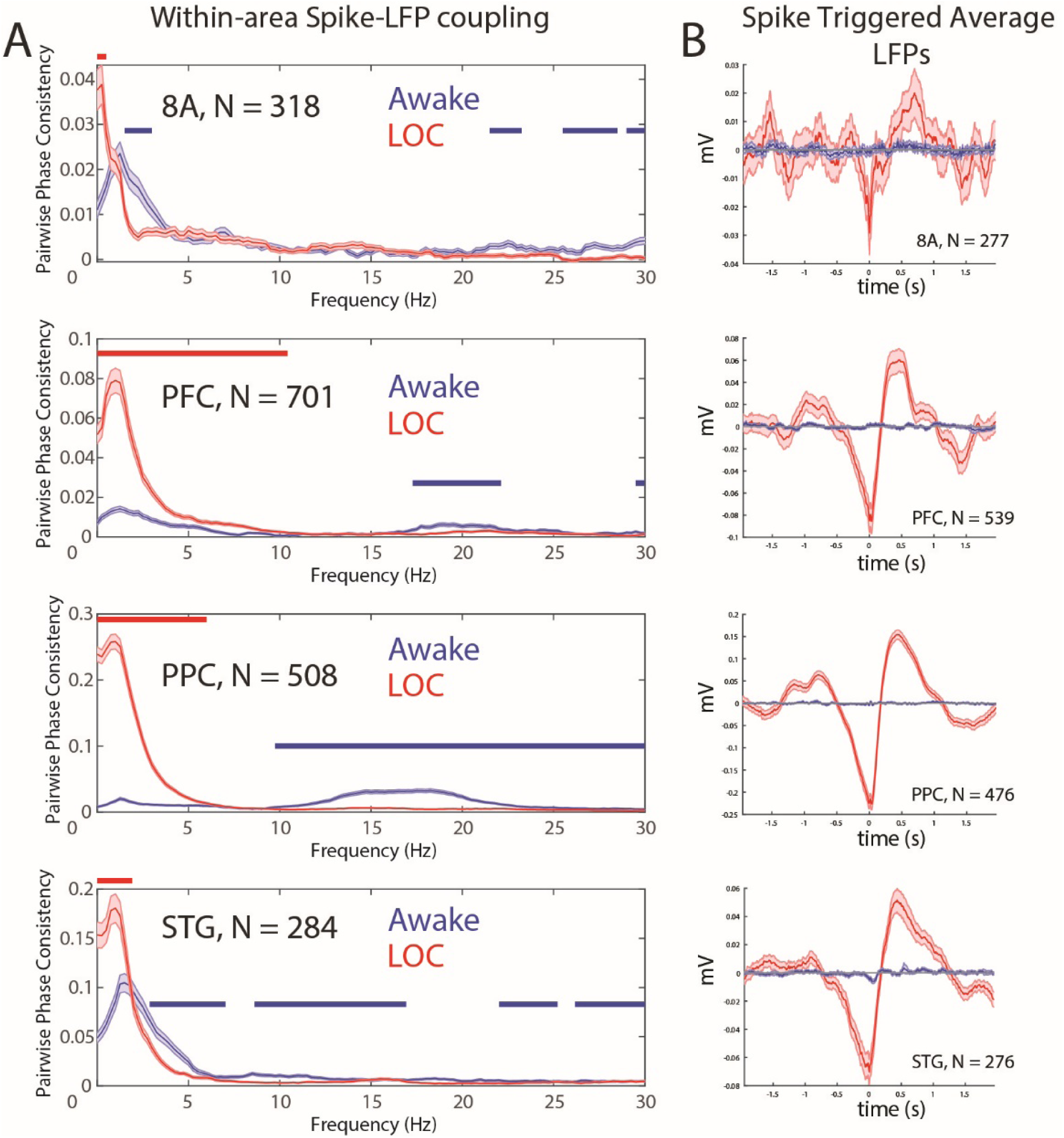
Changes in spike-LFP coupling during awake vs. LOC states. A. Pairwise Phase Consistency of within-area spike-LFP coupling during awake (blue) and post-LOC (red). Spikes and LFPs from the same array but different electrodes. N = number of single neurons. B, Spike-triggered average of LFPs during awake (blue) and post-LOC (red). Error bars indicate +/-SEM. Zero is the time of spiking.

After LOC, spiking was entrained by the increased slow frequency (SF) power, producing Up and Down states of high vs little/no spiking (e.g., Fig 1D) reducing average spike rates (Fig 1E). SF (0.1-1Hz) spike-LFP coupling increased within (Fig 2A) and across (Fig 3A) cortical areas. Spike-triggered averages of the local LFP signal indicated that spikes entrained to the depolarized phases (troughs) of SF oscillations (Fig 2B). Spike-LFP coupling in delta, theta, and beta bands (2-30 Hz) decreased (Fig 3A). SF coupling also increased between cortical spikes and thalamic LFPs (Fig 3B) and between cortical and thalamic LFPs (Fig 3C) while corticothalamic LFP coupling decreased above 4 Hz (Fig 3C). Partial coherence analysis of thalamic LFP on corticocortical LFP coherence showed that the thalamus had distinct frequency-dependent contributions to cortical coherence. Partialing out the contribution of the thalamus decreased within and across-area corticocortical LFP coherence in beta while awake and in SF after LOC (Fig 3B), indicating a thalamic contribution to cortical dynamics.

**Figure 3.**
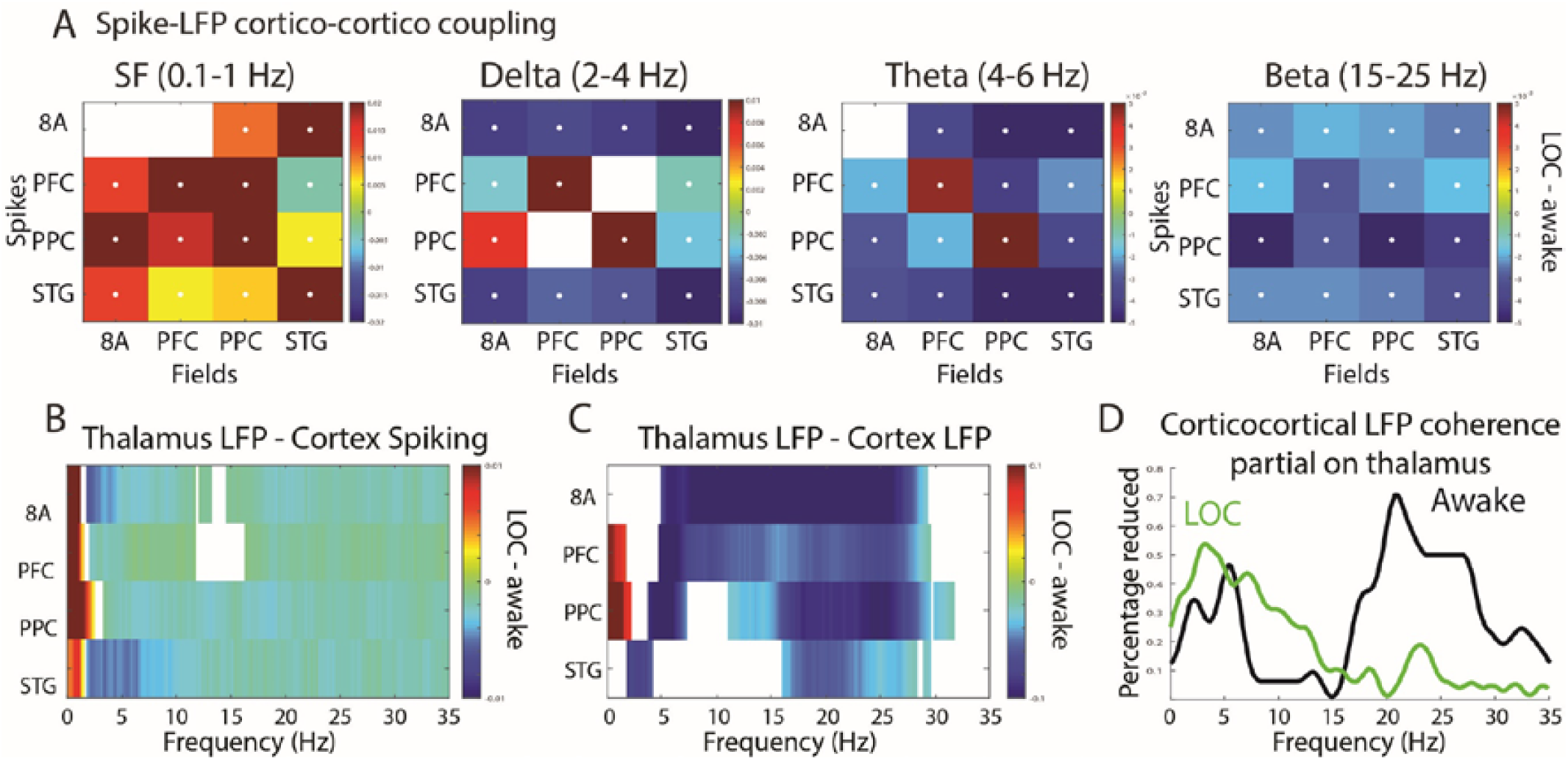
Changes in corticocortical and thalamocortical coherence and partial coherence during awake vs. LOC states. A. Average differences in spike-LFP coupling (LOC minus awake). White asterisk indicates significant difference. X-axis = area providing spikes; Y-axis = area providing the LFPs. No significant differences are masked out. B. Changes in spike-LFP coupling (quantified with PPC) between thalamic LFPs and cortical spikes (LOC minus awake) masked for significance at P<0.01 (see Methods). C. Changes in thalamocortical LFP-LFP coherence (LOC minus awake) masked for significance at P<0.01 (see Methods). D. Percentage of cortico-cortical coherence connections with significantly (P<0.01) reduced coherence due to thalamic partialization; awake (black), post-LOC (green).

Thus, we tested whether we could “wake” the cortex by electrical stimulation of the thalamus. Under the holding dose of propofol, we applied 180 Hz, bipolar stimulation targeting central thalamus (Figure 1A, see Methods)^12^. The animals’ eyes opened (Figure 4A), they began responding to airpuffs (Figure 4B), and pupil diameter and heart rate increased (Figure 4C,D). Stimulation also produced an awake-like cortex, increasing spike rates (Fig 4E), and making them less periodic (Fig 4F). SF LFP power reduced, especially in frontal cortex (8A, and PFC); high-frequency power returned (Figure 4G). Stimulation-induced behavioral and cortical wakefulness persisted 30 sec or more after stimulation (Fig 4A-F) although the decreased SF and increased higher-frequency power could last longer (Fig. 4G). Effective stimulation sites were near or in the ILN (Fig. 1B).

**Figure 4.**
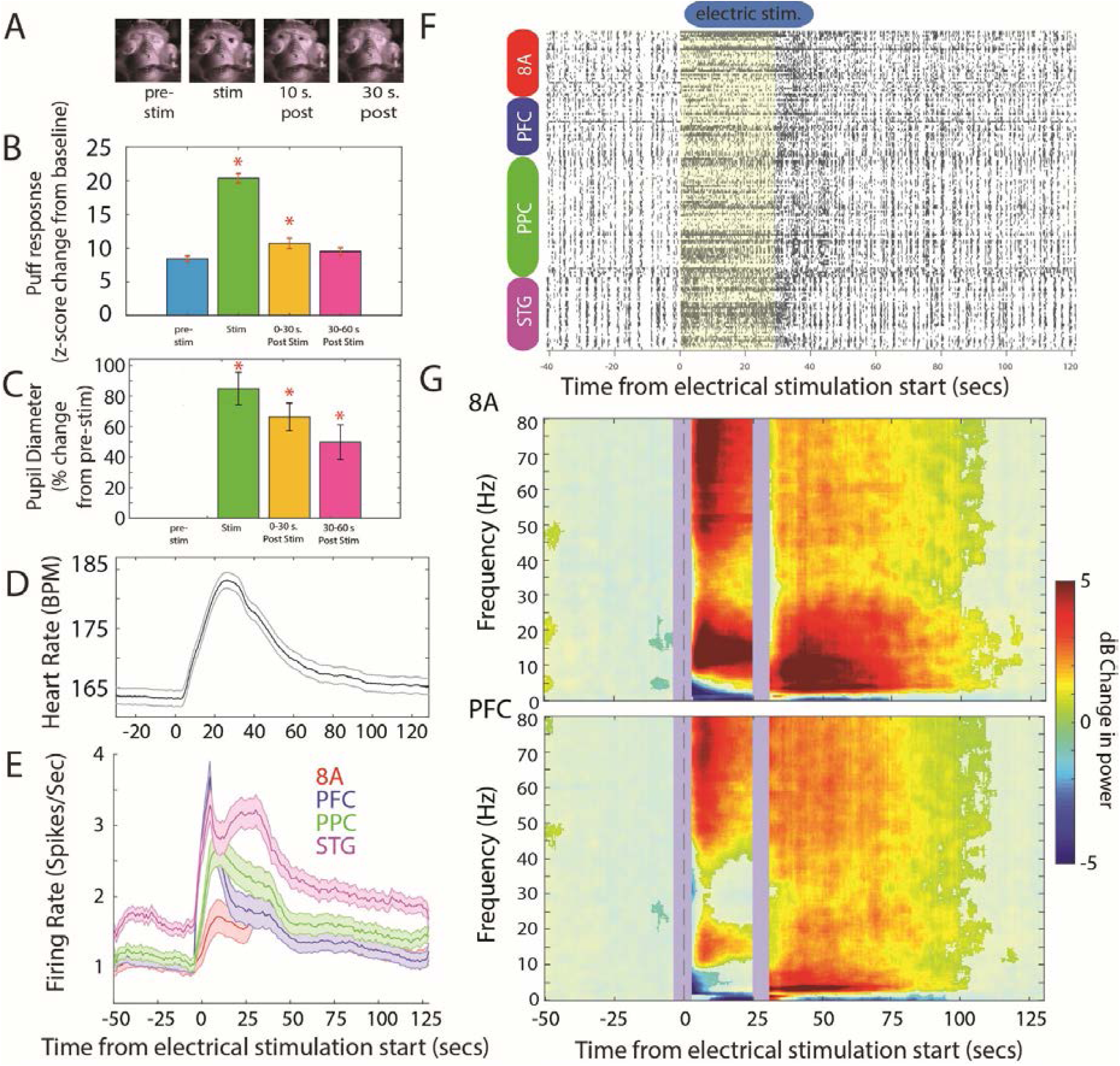
Electrical stimulation of thalamus reverses effects of propofol. A. Representative picture of a monkey pre, during, and post electrical stimulation of the thalamus. B. EMG of response to airpuff to eye, expressed as a z-score change in signal compared to the 200ms pre-airpuff. Error bars are SEM for all figures. Red stars indicate significant changes from pre-stim, p<0.05, t-test. C. Pupil diameter percent change from pre-stim. Note: on some sessions, the monkey’s eyes were propped open, allowing continuous monitoring of pupil size throughout pre/stim/post stim epochs, D, Heart rate (bpm) over time from start of thalamic stimulation (zero). E. Mean spike rates for each cortical area. F. Example of spikes from a single trial with thalamic stimulation. Yellow indicates time of electric stimulation. G, Average change in power for area 8A before and after thalamic stimulation. dB change from 30 seconds preceding stimulation. Significant increases/decreases at P<0.01 (see Methods) are shown with more saturated colors. +/-1.5 seconds at electric stimulation onset and offset are masked due to electric artifact. H. SF (0.1-1.5 Hz) power. I. PPC of SF (0.1-1.5 Hz) spike-LFP coupling. Horizontal bars indicate significant differences (P<0.01).

During unconsciousness, propofol’s GABAA-mediated inhibition enhanced slow-frequency power/coherence between and within areas but decreased coherence in the higher frequencies (>4 Hz) associated with cognition and consciousness^3,6,11,13–17^. Spikes strongly coupled to the depolarized phases of slow-frequency LFPs oscillations, producing Up-states alternating with prolonged Down-states. The thalamus contributed to awake cortical dynamics and the unconscious slow-frequency state. The ILN has diffuse cortical connections posited to mediate the global binding needed for consciousness^18^. Its stimulation has improved responsiveness of minimally conscious patients^12^. We found that it returned to cortex a higher-frequency awake-like state, restoring consciousness. Thus, propofol likely renders unconsciousness by disrupting intracortical communication through enhanced down-states and loss of the higher frequency coherence thought to integrate cortical information^3,4,7,19–24^.

## Acknowledgements

This work was supported by NIMH R01MH115592, NIGM T32GM007753, NIGM P01 118269 and the MIT Picower Institute Innovation Fund. Thanks for Carolyn Wu for histology.

## Supplemental Materials

**Supplemental Figure 1.**
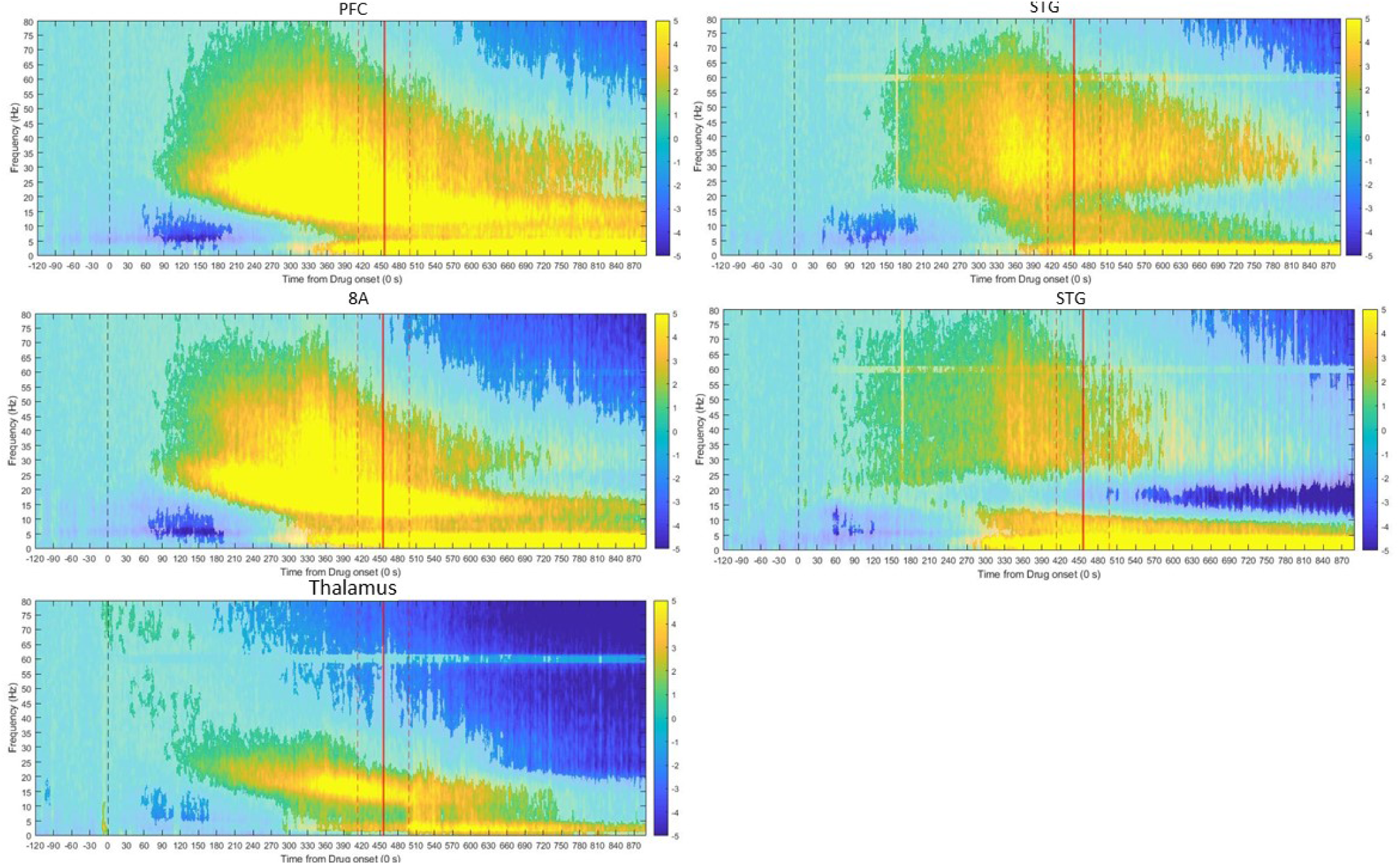
Changes in cortical power for all areas. Same as Figure 2A/B but for all recorded areas.

**Supplemental Figure 2.**
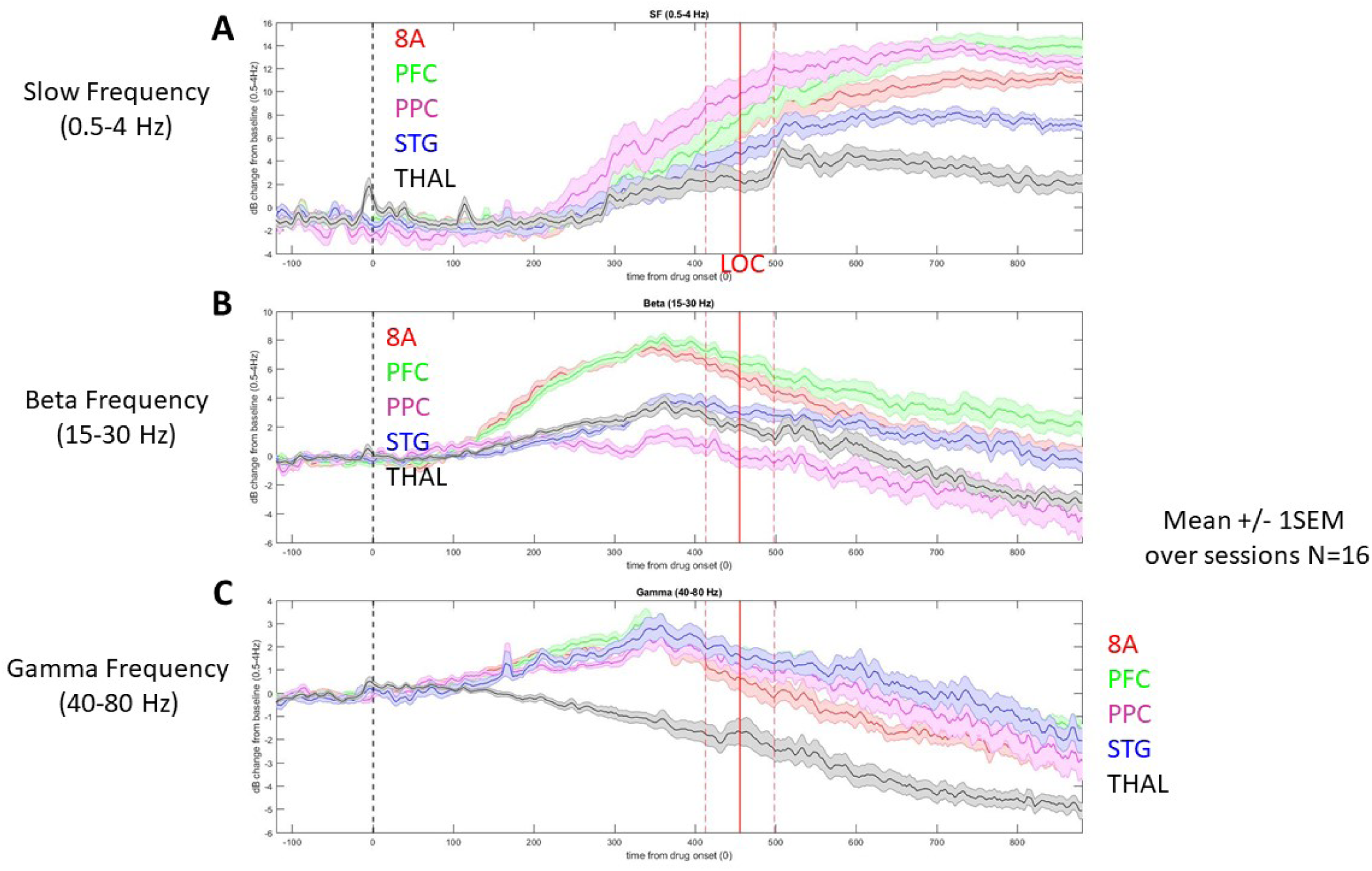
Changes in cortical power per frequency band area. Relative to pre-drug baseline, changes in the average power spectrum are shown for different areas (see legend for color code). A, SF frequency (0.5-4 Hz), B, Beta frequency (15-25 Hz), C, Gamma frequency (40-80 Hz). Mean +/-1SEM

## Methods

Two rhesus macaques (*Macaca mulatta*) aged 14 years (MJ, male, ∼13.0 kg) and 8 years (Monkey LM, female, ∼6.6 kg) participated in these experiments. Both animals were pair-housed on 12-hr day/night cycles and maintained in a temperature-controlled environment (80°F). Each monkey was surgically implanted with a subcutaneous vascular access port (Model CP-6, Norfolk Access Technologies, Skokie, IL) at the cervicothoracic junction of the neck with the catheter tip reaching the termination of the superior vena cava via the external jugular vein.

Monkeys were head-fixed via a titanium headpost and placed in noise isolation chambers with masking white noise (50 dB). Propofol was intravenously infused via a computer-controlled syringe pump (PHD ULTRA 4400, Harvard Apparatus, Holliston, MA). The infusion protocol was stepped such that unconsciousness was induced via a higher rate infusion (285 µg/kg/min for monkey MJ; 580 µg/kg/min for monkey LM) for 15 minutes before dropping to a GA maintenance dose (142.5 µg/kg/min for monkey MJ; 320 µg/kg/min for monkey LM) for an additional 45-90 minutes. Heart rate and oxygen saturation were monitored continuously and recorded throughout all phases of experiments using clinical-grade pulse oximetry (Model 7500, Nonin Medical, Inc., Plymouth, MN). SpO_2_ values were maintained at values above 93% during each recording sessions.

Infrared monitoring tracked facial movements and pupil size (Eyelink 1000 Plus, SR-Research, Ontario, CA) throughout the course of the experiments. Loss of consciousness (LOC) was deemed by the timestamp of the moment of eyes-closing and lack of eyeblink to an airpuff as well as the reduction in suprascapular EMG tone. Recovery of consciousness (ROC) was classified as the timestamp of the first to occur between eyes reopening with spontaneous blinking or regaining of motor activity in the setting of active or ceased drug infusion. To ensure propofol clearance from tissues and physiological recovery, experiments were never repeated on subsequent days or with ketamine anesthesia given within twenty-four hours. All procedures followed the guidelines of the MIT Animal Care and Use Committee and the US National Institutes of Health.

For electrical stimulation of the thalamus, we adapted electrical stimulation parameters previously shown to cause behavioral improvements in coma patients and awake, behaving monkeys ^1,2^. We unilaterally delivered 180 Hz bipolar, biphasic, square wave pulses (1-3.2 milliAmps) between 6-8 contacts on monkey MJ and 6 contacts on monkey LM. Five minutes into the maintenance anesthesia dose (20 minutes from infusion start), 30-second “trials” of electrical stimulation were delivered as the propofol infusion continued. Stimulation trials were separated by 2 minutes intervals, except the 4th and 5th stimulation runs, which were separated by a 5-minute inter-trial interval. These stimulation washout periods sufficiently allowed for reestablishment of the behaviorally-judged anesthetized state (e.g. loss of puff responses). Stimulation of 1 or more mAmp were generally successful in awakening the animal. During a subset of stimulation trials, the eyelids were mechanically propped open to evaluate the changes in pupil size with electrical stimulation. The diameter of the pupil was captured from the infrared eye tracker, with saline flushes applied every 2-3 minutes.

### Neurophysiology

For recordings in cortex, monkeys were chronically implanted with four 8×8 iridium-oxide contact microelectrode arrays (“Utah arrays”, MultiPort: 1.0 mm shank length, Blackrock Microsystems, Salt Lake City, UT), for a total of 256 electrodes distributed in the prefrontal (area 46 ventral and 8A), posterior parietal (area 7A/B), and temporal-auditory (caudal parabelt area STG, Superior Temporal Gyrus) cortices. Specific anatomical targeting utilized structural MRIs of each animal and a macaque reference atlas, as well as visualization of key landmarks on surgical implantation^3^. For Utah array recordings, area 8A and PFC were ground and referenced to a common subdural site. Area STG and PPC also shared a common ground/reference channel which was also subdural. LFPs were recorded at 30 kHz and filtered online via a lowpass 250Hz software filter and downsampled to 1 kHz. Spiking activity was recorded by sampling the raw analog signal at 30 kHz, bandpass filtered from 250Hz-5kHz. Blackrock Cereplex E headstages were utilized for digital recording via a Blackrock Cerebus Digital Acquisition system. Single units were sorted manually offline using principal component analysis with commercially available software (Offline Sorter v4, Plexon Inc., Dallas, TX). All other pre-processing and analyses were performed with Matlab (The Mathworks, Inc, Natick, MA).

For recordings in central thalamus, monkeys were chronically implanted with four 6-8 channel recording/stimulating electrodes (0.5 mm contacts with 0.5 mm intercontact-spacing, NuMed Inc, Hopkington, NY) bilaterally targeting the intralaminar nuclei. A specialized anatomical localization and insertion protocol utilizing serial intraoperative MRIs was developed in order to allow for precise subcortical targeting of the electrodes along the long axis of the central lateral nucleus and extending ventrally to the centromedian and parafasicular nuclei. Custom-reamed carbon PEEK recording chambers were affixed to the skull with acrylic and ceramic screws stereotaxically determined to target the central thalamus. Recording chamber grids with 1mm grid holes were inserted into the chambers, filled with sterile saline, and the monkey’s head was then imaged by 3T MRI. Subsequent confirmation of the appropriate grid holes targeting the thalamic structures of interest, monkeys were generally anesthetized and brought to the operating facility where a small-bore craniotomy (<2mm) was performed at the appropriate grid holes. The grid was replaced in the chamber and the monkey was transferred anesthetized to the imaging facility and administered a gadofosveset trisodium contrast agent to highlight vasculature obstructing the trajectory to thalamus (e.g. thalamostriate vein). In the MRI suite, stylette-cannulae were inserted into the relevant grid holes and lowered several mm into cortex and one set of 0.5 mm resolution images was obtained. Upon confirmation of correct trajectory on MRI, the stylette-cannulae were lowered to their final position, with the tip approximating the thalamus. The stylettes were then removed, and electrodes of marked length lowered to the depth of the cannulae. Following another MRI-based measurement (scan 2) with the electrodes still in the cannulae, the electrodes were lowered to their final positions within the thalamus and reimaged (scan 3). Upon final assessment of correct localization, the probes were fixed in place and the chamber sealed with acrylic. Histological staining with acetylcholinesterase was used to confirm exact electrode contact locations within and outside the central thalamus of both monkeys.

Thalamic recordings were referenced to the monkey’s titanium headpost. In offline analysis, thalamic sites were re-referenced to a nearest-neighbor bipolar montage prior to further analysis. LFPs were recorded via a separate analog front end amplifier and an additional identical digital acquisition system, synchronized to the digital acquisition utilized for cortical recordings. LFPs were similarly recorded at 30 kHz and filtered online via a lowpass 250Hz software filter and downsampled to 1 kHz.

At the end of the recording sessions, the animals were sacrificed for histological confirmation of thalamic sites, as previously described^4^. Briefly, monkeys were given a lethal dose of sodium pentobarbital. When they became areflexive, they were perfused transcardially with PBS, followed first by a cold solution of 4% paraformaldehyde and next by a mixed solution of 4% paraformaldehyde and 10% sucrose. Blocks of brain and spinal cord were removed and stored overnight in 30% sucrose at 5°C before cutting. Sections were processed for acetylcholinesterase. Anatomical localization of electrodes was determined by histological examination of brain tissue. It was not necessary to create electrolytic lesions prior to histology, because the thalamic electrodes were wide enough (0.5mm diameter) to be unambiguously identified in the anatomical sections.

Electrical stimulation generally produced artifacts that were highly correlated across channels. We removed these artifacts using ZCA-whitening^5^. ZCA whitening is the linear whitening transformation that minimizes the mean squared error between the original and whitened signals. First, we applied a fourth-order Butterworth band-pass filter to the raw, 30 kHz recorded signals between 300 and 4500 Hz. Next, we estimated the cross-channel covariance matrix Q from the filtered signals recorded during DBS stimulation. Finally, we multiplied the matrix of filtered signals by Q-1/2 to yield the whitened signals. For computational efficiency, we computed the cross-channel covariance matrix using a subset of 100,000 randomly selected time points. From these whitened signals, we manually set voltage thresholds to capture high-frequency spikes and sorted these units using principal component analysis in commercially-available software (Offline Sorter v4, Plexon Inc., Dallas, TX).

### Data Analysis

Local Field Potentials (LFPs) were analyzed using the Chronux ^6^ for spectrograms and the Fieldtrip toolbox for other analyses^7^. For each channel on each array or thalamic probe, we computed a time-frequency decomposition using the Chronux analysis toolbox. We used the multitaper method with 1 Hz spectral smoothing to estimate power in 1 Hz intervals from 1 to 80 Hz and at non-overlapping 0.5 second time intervals. We calculated power from 120 seconds prior to propofol onset to 1180 seconds post onset. Each window of analysis consisted of 4 seconds of data, giving a frequency resolution of 0.25 Hz. We then averaged across all electrodes in a given area, to get a clean estimate of each area’s frequency response profile, per session. All thalamus leads were combined for power/coherence analysis.

We calculated the change in power between drug and pre-drug baseline in decibel (dB) units. In other words, we applied the following transformation to the raw power values:

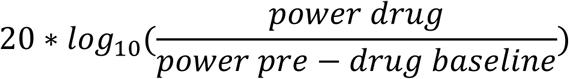

We took the pre-drug baseline to be the 2 minutes immediately before drug administration, and the drug period to be every second after drug administration. For each area, we then assessed whether there was significant power modulation by using a non-parametric cluster-based randomization test^8^. For each session, we used the null hypothesis that power in the baseline and power in the drug period were the same. To this end, we randomly exchanged baseline-transformed time-frequency power estimates between the baseline and drug windows. We extracted the largest cluster (continuous tiles in time-frequency space) to pass a first level significance threshold, by applying a t-test and thresholding all significant bins p<0.01, uncorrected. We performed this randomization 1,000 times. The empirically observed clusters were compared to this randomization distribution to assess significance at p<0.01, adjusted for multiple comparisons across sessions.

To calculate the effect of electrical thalamic stimulation on cortical power, we applied a similar transformation, only this time the baseline was the 30 seconds of data immediately preceding each trial of stimulation:

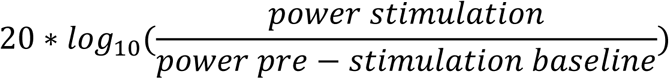

We then repeated the same test outlined above, only now randomizing bins before/during/after stimulation onset to create the null hypothesis. There were stimulation onset and offset artifacts. We removed the times around onset/offset +/-3 seconds prior to performing this randomization test. As a result, these artifact times are omitted from the analysis and figure.

### Methods, spike-LFP coherence and partial coherence analysis and statistics

For each area, we calculated the average LFP across the 64 channel array. We then used the Pairwise Phase Consistency (PPC) metric to calculate coupling between spikes and LFPs. This was implemented in Fieldtrip in the ft_functionalconnectivity function^9^, setting method to ‘ppc’. Note that the PPC metric is not biased by the spike count^10,11^, thus enabling comparison between awake and loss of consciousness (LOC) and recovery of consciousness (ROC) states without a statistical confound. To determine whether spike-field coupling (quantified with the PPC metric) was significantly different between awake and LOC conditions, we first quantified the average PPC across frequencies separately for both conditions (awake vs LOC), taking all available spikes and fields for each condition separately. Thus, each single unit provided 1 PPC measurement from LOC and 1 PPC measurement from the awake state. We then computed significant differences by randomly shuffling the label, re-computing the difference in PPC in the shuffled data, and performing cluster-based randomization testing to determine significant at a multiple comparisons corrected value of p<0.01.

To determine whether the thalamus mediated corticocortical coherence, we first computed coherence between LFPs using Fieldtrip, using the function ft_connectivityanalysis with method ‘coh’. We then computed partial coherence for the same corticocortical pairs, but with the thalamus partialed out. We used the partial coherence method^12^, implemented in ft_connectivityanalysis with method ‘pcoh’. Therefore, each recording session and each corticocortical combination provided two coherence values: the raw coherence and the partial coherence. Using cluster-based randomization testing, we determined whether partial coherence significantly reduced the regular coherence. To do this, we randomized whether an interaction was partial or non-partial across sessions, thus realizing the null hypothesis that partial and non-partial coherence are not different. We determined significant at p<0.01, corrected for multiple comparisons. If a frequency bin/cortico-cortical interaction was significantly reduced by partialization given this test, we included it in the numerator for Figure 3D, to determine the percentage of cortico-cortical connections that were significantly reduced by thalamic partialization.

